# Network analysis of the hominin origin of Herpes Simplex virus 2 from fossil data

**DOI:** 10.1101/105007

**Authors:** Simon Underdown, Krishna Kumar, Charlotte Houldcroft

**Affiliations:** Human Origins and Palaeoenvironmental Research Group (HOPE), Department of Anthropology & Geography, Oxford Brookes University, Oxford, OX3 0BP, UK; Leverhulme Centre for Human Evolutionary Studies, University of Cambridge, Henry Wellcome Building, Fitzwilliam Street, Cambridge, CB2 1QH, UK; Computational Geomechanics, Cambridge University Engineering Department, Trumpington Street, Cambridge, CB2 1PZ, UK; Division of Biological Anthropology, Department of Archaeology & Anthropology, University of Cambridge, Cambridge, CB2 3QG, UK; McDonald Institute for Archaeological Research, University of Cambridge, Downing Street, Cambridge, CB2 3ER, UK

**Keywords:** network analysis, human evolution, infectious disease, epidemiology, archaeology

## Abstract

Herpes simplex virus 2 is a human herpesvirus found worldwide that causes genital lesions and more rarely causes encephalitis. This pathogen is most common in Africa, and particularly in central and east Africa, an area of particular significance for the evolution of modern humans. Unlike HSV1, HSV2 has not simply co-speciated with humans from their last common ancestor with primates. HSV2 jumped the species barrier between 1.4 and 3 MYA, most likely through intermediate but unknown hominin species.

In this paper, we use probability-based network analysis to determine the most probable transmission route between intermediate hosts of HSV2, from the ancestors of chimpanzees to the ancestors of modern humans, using paleo-environmental data on the distribution of African tropical rainforest over the last 3 million years and data on the age and distribution of fossil species of hominin present in Africa between 1.4 and 3 MYA. Our model identifies *Paranthropus bosses* as the most likely intermediate host of HSV2, while *Homo habilis* may also have played a role in the initial transmission of HSV2 from the ancestors of chimpanzees to *P boisei.*

## Introduction

Herpes simplex virus 2 (HSV2) is a sexually transmitted human pathogen that causes genital lesions and, rarely, encephalitis (eg Tang et al. (2003)), and is associated with increased risk of HIV acquisition (Freeman et al., 2006). After primary infection, the virus adopts a life cycle of latency punctuated by periods of lytic replication when new hosts can be infected through genital contact. The virus is related to the human oral pathogen herpes simplex virus 1 (HSV1). Both HSV1 and HSV2 are alphaherpesviruses, which are found in many primates (Wertheim et al., 2014). HSV1 primarily causes infection, and sporadic lesions, within the oral cavity and establishes latency in the trigeminal ganglia, while HSV-2 is associated with infection of the genitalia and surrounding skin, and establishes latency in the sacral ganglia (Whitley, Kimberlin, and Prober, 2007). Both simplex viruses can infect either body cavity, although there is viral shedding data from co-infected individuals to suggest that HSV1 is a more successful oral and HSV2 a more successful genital pathogen (Kim et al., 2006). The genetic basis of this difference in tropism isn’t fully understood: some studies of recombinant HSV1 × HSV2 strains have highlighted the importance of the latency-associated transcript (LAT) in how successfully the two simplex viruses reactivate from latency within different nerve types (Bertke, Patel, and Krause, 2007). HSV1 and 2 also differ in the length of the glycoprotein G (US4) open reading frame, which may play a role in tropism (Baines and Pellett, 2007).

HSV2 was originally thought to have co-speciated with humans when our lineage diverged from that of the ancestors of chimpanzees and bonobos (anc-chimps). Comparisons of the HSV1, HSV2 and chimpanzee herpes virus 1 (ChHV1) genomes (Tang et al., 2003) suggest that HSV2 is more closely related to ChHV1 than HSV1 (Wertheim et al., 2014). This analysis also found that ChHV1 and HSV2 diverged from one another between 1.4 and 3 MYA, and the authors inferred that an unknown hominin (or hominins) was infected with HSV2 before it switched host to the ancestors of modern humans.

We hypothesise that by combining fossil data on when and where different hominin species were likely to be present in Africa, the geographical range of modern chimpanzees and bonobos, and the reconstructed distribution of tropical rainforest habitat as a proxy for the past range of the chimpanzee/bonobo ancestor (anc-chimps) it will be possible to develop a model to statistically infer the species that facilitated the host-switch of HSV2 in the modern human lineage defined here as beginning with *Homo erectus* (Anton et al., 2016).

Before *Homo erectus* it is unclear which hominins are the direct ancestors of anatomically modern humans. However, once HSV2 has reached *Homo erectus,* no further host-switches are required for HSV2 to be considered present in the ancestors of modern humans.

HSV2 is found in all living populations (Looker et al., 2015), is accepted as having an African origin (Koelle et al., 2017; Burrel, Boutolleau, et al., 2017), and has patterns of genetic divergence consistent with having diverged with human populations as they spread out of Africa (Koelle et al., 2017). Current genetic, archaeological and fossil evidence suggests 100 KYA as a plausible (although not universally accepted) earliest date for anatomically modern humans (AMH) to have left Africa Mirazon Lahr et al., 2016. We therefore infer that HSV2 must have been in the population of AMH before they left Africa, in order to take it with them when migrating to the rest of the world.

## Methods

### HSV2 prevalence data, hominin fossil data and chimpanzee and tropical rainforest geographic range data

HSV2 is most closely related to ChHV1 which infected the ancestors of modern chimpanzees. The GIS data on the range of modern *Pan troglodytes* and *Pan paniscus,* provided by the IUCN Red List (Oates et al., 2008), is shown in fig. 3a. Only one ancestral chimpanzee fossil, dated to c. 500 KYA, is currently known (McBrearty and Jablonski, 2005). This means that the habitat range of anc-chimps is not directly measurable from the fossil record but is inferred to be a larger geographical range. Therefore, we have used the paleo-tropical rainforest range during the period of 1.4 - 3 MYA Koppen-Geiger climate classification dataset Peel, Finlayson, and McMahon, 2007 as a proxy for the ancient range of chimpanzees and combined this with data based on modern great ape distribution patterns and range size estimates (Myers Thompson, 2003), and is shown in fig. 3b.

**Figure 3.**
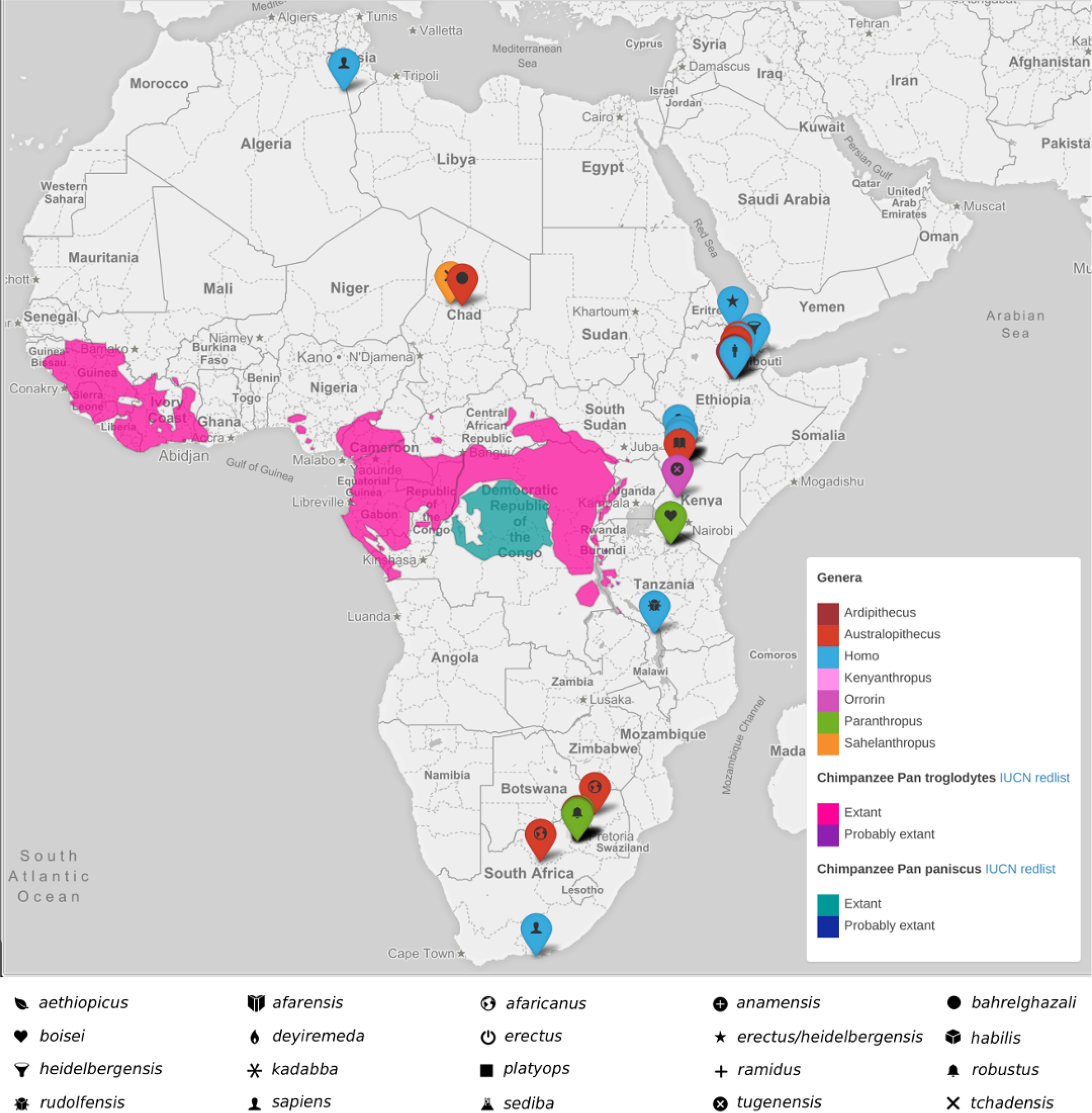
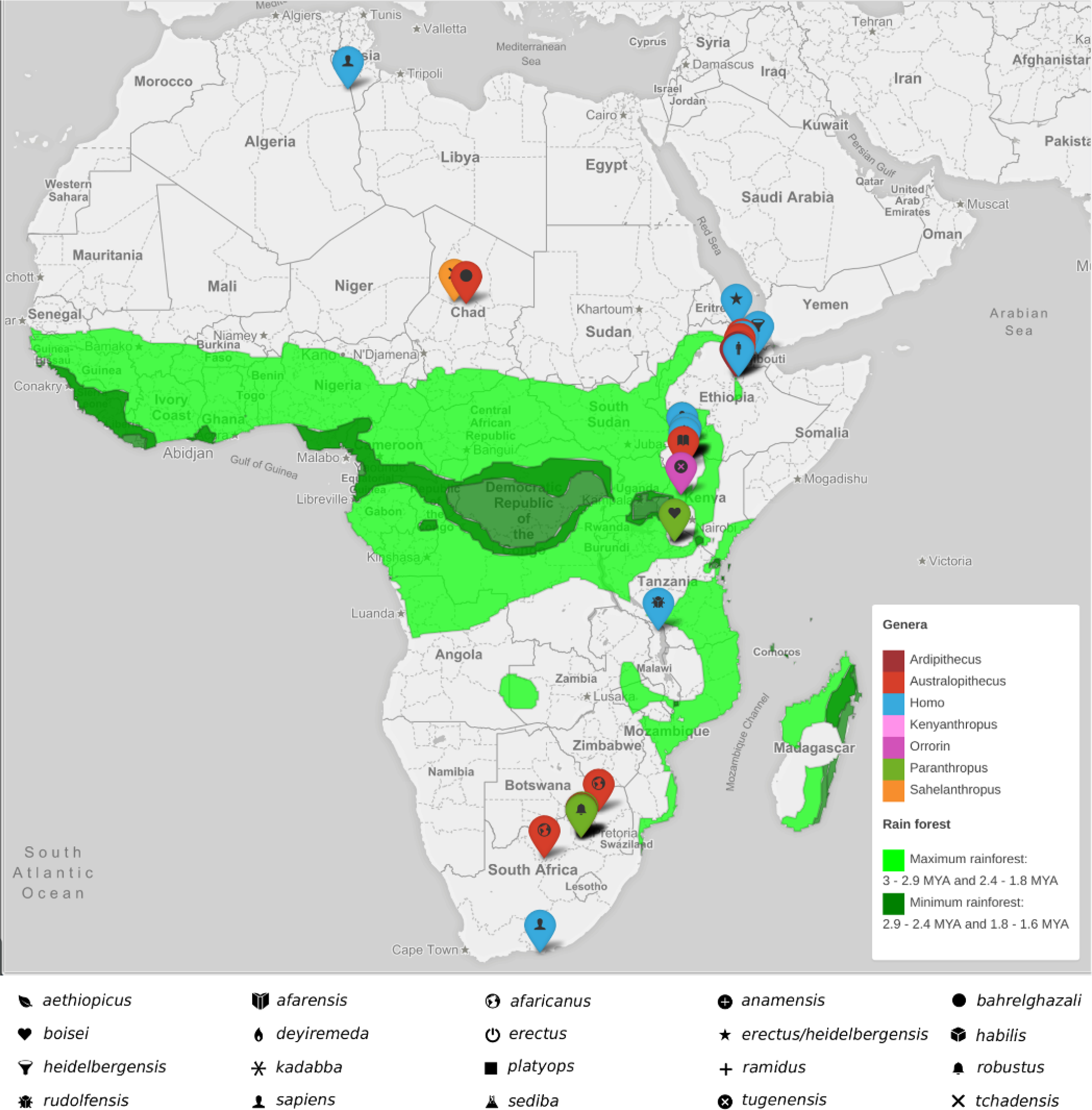
Location of hominin fossils, extant chimpanzees, and ancient minimum and maximum rainforest distributions. (a) Distribution of extant chimpanzee (Pan troglodytes) and bonobo (Pan paniscus) populations [IUCN redlist http://maps.iucnredlist.org/map.html?id=15933, http://maps.iucnredlist.org/map.html?id=15932]. The locations of hominin fossils [supplementary table 1] are shown with markers. The colour of the marker indicates the hominin genus; the symbol represents the species. (b) Location of hominin fossils relative to ancient minimum and maximum rainforest distributions (Peel, Finlayson, and McMahon, 2007). These figure are available interactively: https://wadhamite.github.io/hsv-mapping.

To identify which species could have been involved in the host-switch of HSV2 from the ancestors of chimpanzees to the ancestors of modern humans, we collated spatio-temporal data on African hominin fossil species extant between 100 KYA and 3 MYA. Latitude and longitude of site location was used to provide a spatial data point for each species. Temporally, published fossil dating evidence was used to provide a first appearance datum (FAD) and last appearance datum (LAD) for each species (see supplementary Table 1).

Data on the prevalence of herpes simplex virus 2 between 2000 and 2015 CE was taken from the supplementary materials of (Looker et al., 2015), and plotted to demonstrate the distribution of HSV2 across Africa (Supplementary Figure A. 1).

### Network analysis

To establish the most probable transmission route of HSV2 from anc-chimps to modern humans, it is important to identify potential species that may have been intermediary hosts. Initially, all hominins in Africa with temporal ranges within the 1.4 - 3 MYA confidence window (Wertheim et al., 2014) of the chimp-hominin transmission were identified (see Supplementary Table 1). Their distance to ancient tropical rainforest was calculated, and only those species whose fossil remains were found within 400 km of tropical rainforest were considered as putative species for initial ancestral-chimp-hominin HSV2 transmission. This reflects the distance that would have been covered by hominins employing three possible strategies: (a) a broadly omnivorous subsistence strategy based on scavenging, (b) hunting using carnivore and herbivore movement patterns and (c) modern hunter-gatherer range sizes (Robert A. Foley, 1978; Grant, Chapman, and Richardson, 1992). A matrix of spatio-temporal distances was then calculated to map the distances between the nearest neighbours of each species and also to calculate the temporal overlap between species using the fossil record. Fossils from four genera *(Ardipithecus, Kenyanthropus, Orrorin* and *Sahelanthropus)* were excluded from the analysis on the basis that there is no fossil evidence that they persisted after 3 MYA.

A network of possible transmission routes of HSV2 from anc-chimps to humans through different hominins was developed as a directed acyclic graph (DAG) *G* = (V, *E*) comprising of a set of nodes (V), representing potential intermediary hosts, and edges (E) connecting the nodes, which represents the direction of transmission between species (See Figures 1 and 2). The DAG comprised of the anc-chimps as the start node, *H. erectus* as the target node, and other potential species forming secondary nodes in the graph. The edges are typically weighted; in this analysis, the weights are based on the inverse probability of transmission.

If HSV2 was transmitted to *Homo erectus,* no further cross-species transmission event is needed to explain the infection of modern humans. Simple vertical mother-to-child or horizontal (sexual) transmission of the virus through the genus *Homo* from this point would be sufficient as the ancestor-descendent path from *Homo erectus* to *Homo sapiens* is relatively secure (Maslin, Shultz, and Trauth, 2015).

#### Bayesian inference

A Bayesian network or a belief network is a graphical structure that allows us to represent and reason about an uncertain domain. In a Bayesian network, the species (nodes) are variables, and the transmission routes (edges) represent direct links between species. The process of conditioning (also called probability propagation or inference) is performed via a “flow of decisions” through the network, which involves computing the posterior probability distribution for a set of query nodes, given values for some evidence (or observation) nodes. An important consideration in all Bayesian-based methods is the choice of a prior. An empirical Bayesian method that estimates the likelihood of HSV2 infection using a prior beta distribution is adopted, which is an approximation to a hierarchical Bayesian approach (Farine and Strandburg-Peshkin, 2015; Murphy, 2012).

Bayesian networks provide full representations of probability distributions over their variables, which allows us to infer upon any subset of variables. A Bayesian network is created using the DAG described above. Each node on the graph represents a potential intermediary host that has a probability of transmitting HSV2. The probability of infection transmission is represented as a beta distribution, with shape parameters *alpha* representing the time period of the species in 100,000 years and *beta* representing the distance to the neighbour in kilometres. A *Conditional Probability Table* (CPT) is generated for all possible combinations of a dichotomous outcome for each variable (HSV2 infection is true or false). The *“Combined Inference”* approach is adopted to evaluate the probability of intermediary hosts transmitting HSV2, by conditioning the ancestral-chimpanzee and *Homo erectus* nodes for the presence of HSV2 (Korb and Ann E. Nicholson, 2003). The Bayesian inference analysis was performed using the AI space decision network tool (Poole and Mackworth, 2010).

#### Optimal path traversal

A* is an informed search algorithm that searches through all possible paths to the target that yields the smallest cost (Hart, Nilsson, and Raphael, 1968). This is done by combining information by favouring vertices that are close to the starting point and to the target. At each time-step the A* algorithm selects the path at a given vertex *n* that has the lowest *f (n) = g(n) + h(n),* where *g(n)* represents the exact cost of the path from the starting point to any vertex n, and *h(n)* represents the heuristic estimated cost from vertex *n* to the goal. A* balances the two as it moves from the starting node to the target node. The most probable transmission route is evaluated by minimising the traversal costs based on the edge cost and nodal heuristics of the transmission network graph. A Python script with the *NetworkX* package was used to implement the A* path-finding algorithm on the DAGs to identify the most probable HSV2 transmission (optimal path) route.

anc-chimps and Homo erectus formed the start node and the target node of the DAG. The edges represent the direction of HSV2 transmission and are weighted based on the inverse probability of transmission between the species (nodes). The weights (edge costs) are determined based on two probability models: Infection Prevalence (HSV2-IP) and Infection Transmission (HSV2-IT) based on the temporal and geographic range of the species. More details about the probability models are discussed in the next section. A Conditional Probability Table (CPT) is used to determine the nodal heuristics.

#### HSV2-Infection Prevalence (HSV2-IP) model

The Infection Prevalence model is a local model of probability, which assumes each transmission route and species to be independent. The model uses a beta distribution to determine the probability of a species transmitting/infected-by HSV2 based on the proximity of the species to the rainforest habitat and the duration (difference between the first and the last appearance datum) of the species. Both the probability of transmission and infection are described using beta distributions. The probability density function for a beta distribution is defined as:

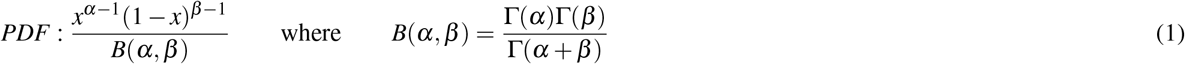

Where, Γ defines a gamma distribution function, *α* is the time period of existence of the species in 100,000 years and β is the spatial distance of the fossil from the rainforest in kilometres. The probability of transmission through a particular route is determined as the combined probability of the species (nodes) forming the edge. For example, the probability of transmission route from ancestral-chimps (*P(A)*) to *P boisei (P(B))* is the combined probability of both these species *P(A * B)*. The edge costs were estimated by Monte-Carlo simulations of the combined probabilities of two species (nodes), forming the edge, by sampling from their respective beta distributions.

#### Stochastic modelling of infection transmission

Infection transmission in epidemiology can be predicted using mathematical models: a popular approach is the Susceptible - Infection - Recovered (SIR) model (Chen et al., 2008). In the stochastic version of the SIR model, the continuous variables are replaced by discrete numbers, and the process rates are replaced by process probabilities. At time *t* the probability that a new susceptible host is infected is modelled as an exponential distribution, which is epidemiologically incorrect for most diseases (Wearing, Rohani, and Keeling, 2005; Bailey, 1975; Sartwell et al., 1950), i.e., the rate of leaving the exposed is independent of the time spent on the host. Wearing, Rohani, and Keeling (2005) suggested a more realistic distribution of latent and infectious periods, with a stronger central tendency:

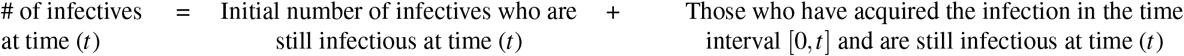

More realistic distributions can be obtained by choosing probability density function of the infectious period, p(t) to be a gamma probability density function (Blythe and Anderson, 1988; Lloyd, 2001).

#### Infection Transmission (HSV2-IT) model

The proposed Infection Transmission model considers the history of transmission, and the probability of routes (edges) and species (nodes) are dependant on the parent nodes and routes. The model utilises the temporal overlap between hominin species and their geographic proximity to one another to determine the probability of a transmission route, using a gamma distribution. The probability density function of a gamma distribution is given as:

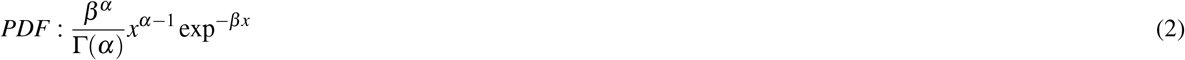

in terms of shape α and rate *β*. The shape parameter α is defined as the ratio of the time period in 1000 years / distance in kilometres and the rate parameter *β* is defined as the normalised time period *Y* /x, where × is the time period of the species in 1000 years and Y is the time period of anc-chimps. Monte-Carlo simulations were performed to evaluate the conditional probability of transmission between species, as mutually exclusive events, considering the parent nodes by sampling from the respective probability distributions. For example, the probability of *P. bolsei* transmitting HSV2 to *H. erectus* depends on the probabilities of *P Boisei* being infected by the anc-chimps and/or *H. habilis* and/or *H. rudolfensis.*

### Sensitivity analysis

Sensitivity analysis is performed to identify the impact of the probability distribution models in estimating the edge costs and in turn the optimal route of transmission. A variance-based global sensitivity analysis was performed using the Saltelli method, a variation of the Sobol technique, to evaluate the sensitivity indices of the transmission model (Sobol/, 2001; Saltelli et al., 2010). The sensitivity of each input is represented by a numeric value, called the sensitivity index. These indices are used to estimate the influence of individual variables or groups of variables on the model output, by decomposing the variance of the model output into fractions. A 10% variance in the model inputs was assumed. A parametric space of edge costs was generated for the Sobol analysis by sampling 50,000 times from the probabilistic distribution for each edge. This generated a total of 2.3 million analyses for the sensitivity analysis for each transmission model. The first-order, second-order and the total-order indices were measured using a Python script with the *SALib* package.

## Results

The results from the Bayesian inference and the optimal path traversal using the Infection Prevalence and Infection Transmission models are presented below.

### Bayesian inference

By conditioning anc-chimps and H. erectus for the presence of HSV2, the combined Bayesian inference of the DAG reveals *A. afarensis* (90%), *H. habilis* (67.8%), *P boisei* (50%), and *H. rudolfensis* (55.8%) to be the likely intermediary hosts for the transmission of HSV2.

### Optimal path traversal

The optimal traversal route of the DAG for HSV2 transmission was modelled using the Infection Prevalence and the Infection Transmission models. The A* algorithm using the weighted DAGs from both models identified *Paranthropus boisei* as the most probable intermediary hosts that transmitted HSV2 from anc-chimps to the ancestors of modern humans (see fig. 1). In addition to the direct transmission of HSV2 from anc-chimps to *P. boisei*, the Infection Transmission model also identified Homo habilis as an intermediary host that transmitted HSV2 to *Paranthropus boisei,* which subsequently transmitted to ancestors modern humans(see fig. 2). Table 1 shows the probable HSV2 intermediary routes and their rankings based on the edge weights from the probabilistic models.

**Figure 1.**
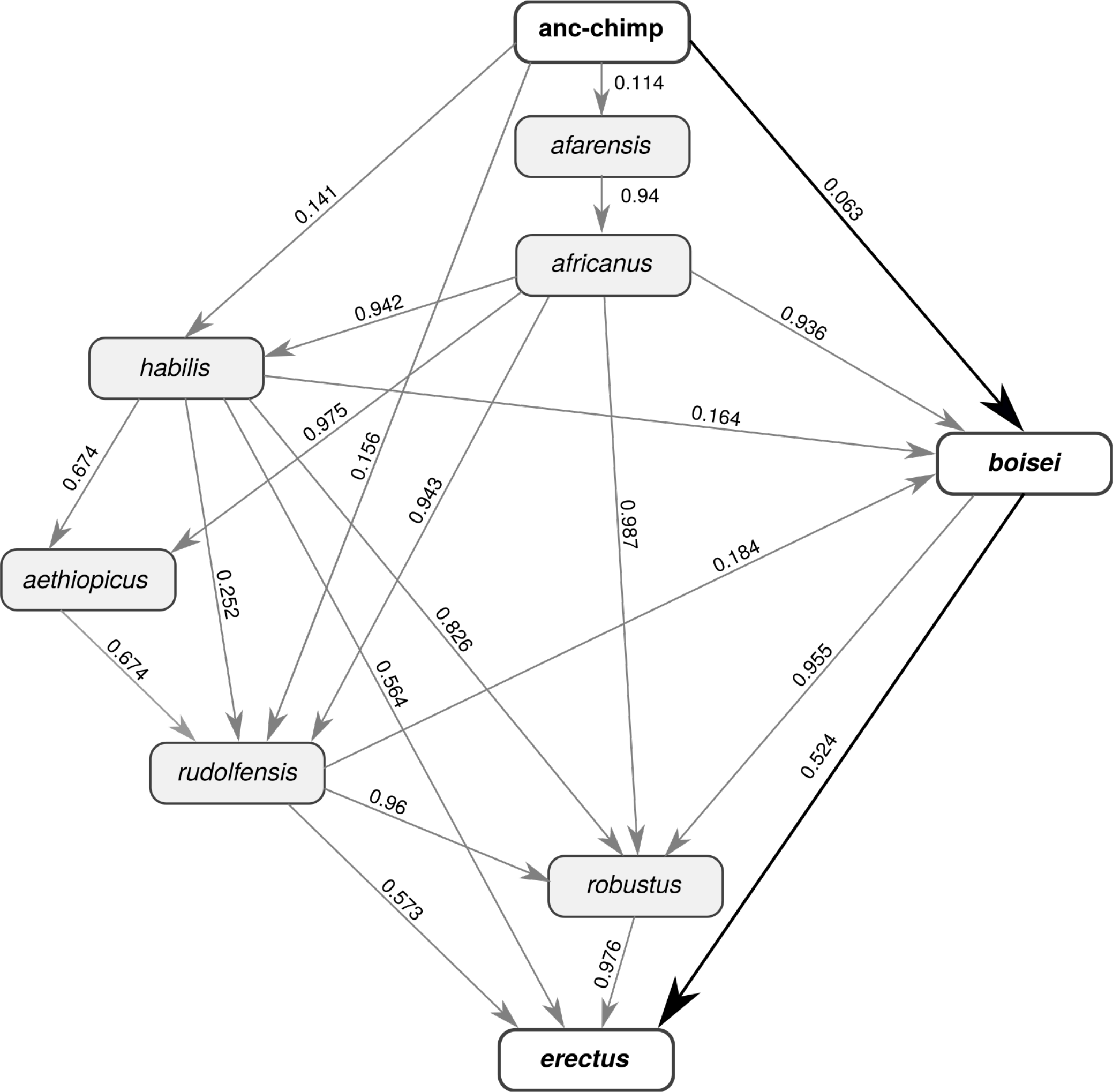
the A* shortest path route(s) for the Infection Prevalence model. The lines with arrows are the possible transmission paths. The values on the lines are the edge costs (inverse probability of transmission). This model predicts that the host switch of HSV2 occurred through the route ancestral-chimps, P. boisei and H. erectus. The route remained unchanged in the sensitivity analysis.

**Figure 2.**
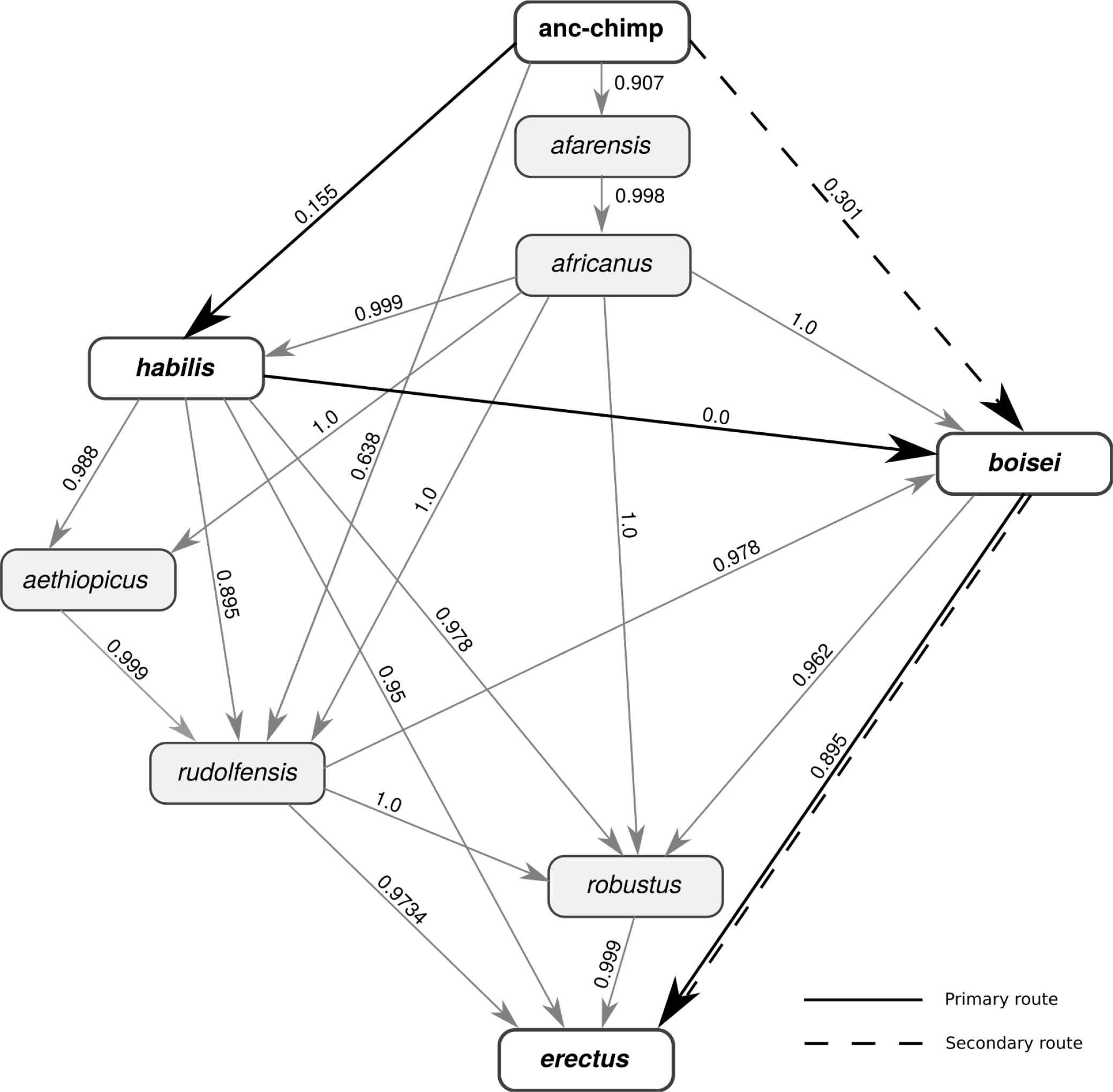
the A* shortest route for the Infection Transmission model. The lines with arrows are the possible transmission paths. The values on the lines are the edge costs (inverse probability of transmission). This model predicts that the host switch of HSV2 occurred through the primary route of ancestral-chimps, *H. Habilis, P boisei* and *H. erectus.* Sensitivity analysis revealed the primary route occurred 60% as opposed to the secondary route of ancestral-chimps, *P boisei* and *H. erectus* (which occurred the remaining 40%).

**Table 1.** Probable HSV2 transmission routes.

| Disease Transmission Route | HSV2-IP route ranking<br>(normalised path cost) | HSV2-IT route ranking<br>(normalised path cost) |
| --- | --- | --- |
| anc-chimp ⇒ *P. boisei* ⇒ *H. erectus* | **1 (1.0)** | 3 (1.14) |
| anc-chimp ⇒ *H. habilis* ⇒ *P. boisei* ⇒ *H. erectus* | 4 (1.13) | **1 (1.0)** |
| anc-chimp ⇒ *H. habilis* ⇒ *H. erectus* | 2 (1.20) | 2 (1.05) |
| anc-chimp ⇒ *H. rudolfensis* ⇒ *H. erectus* | 3 (1.24) | 4 (1.54) |

\* The most probable route has a cost of 1. The path costs are normalised to the shortest path.

### Sensitivity analysis

We performed sensitivity analysis allowing the input parameters of the probabilistic distributions for both the models to vary by 10%. A total of 2.3 million analyses were performed for each model by sampling 50,000 times from the respective probability distributions for each edge.

#### HSV2-IP model

The input parameters to this model: distance to the rainforest and the overlap in the time-period was allowed to vary by 10%. Sensitivity analysis of HSV2-IP model always predicted the transmission route as ‘anc-chimps → *P. boisei → H. erectus’.* The average path cost was 0.587, and it varied between 0.526 and 0.649. The distribution of path costs obtained from the sensitivity analysis is shown in fig. 4. The total-order and first-order Sobol indices of the ‘anc-chimp → *boisei’* path is 4.53E-2 and 4.53E-2, respectively. The ‘P. *boisei → H. erectus’* path has a total index of 9.55E-1 and a first-order index of 9.55E-1. All other paths had negligible or zero index values. When the critical path through *P. bosei* is not available, the transmission happened directly from anc-chimps to *H. erectus* through *H. habilis* (normalised path costs 1.08 - 1.32 of the critical path cost) for 67% of the cases and directly through *H. rudolfensis* (normalised path costs of 1.11 - 1.32 of the critical path cost) for the remaining 33%.

**Figure 4.**
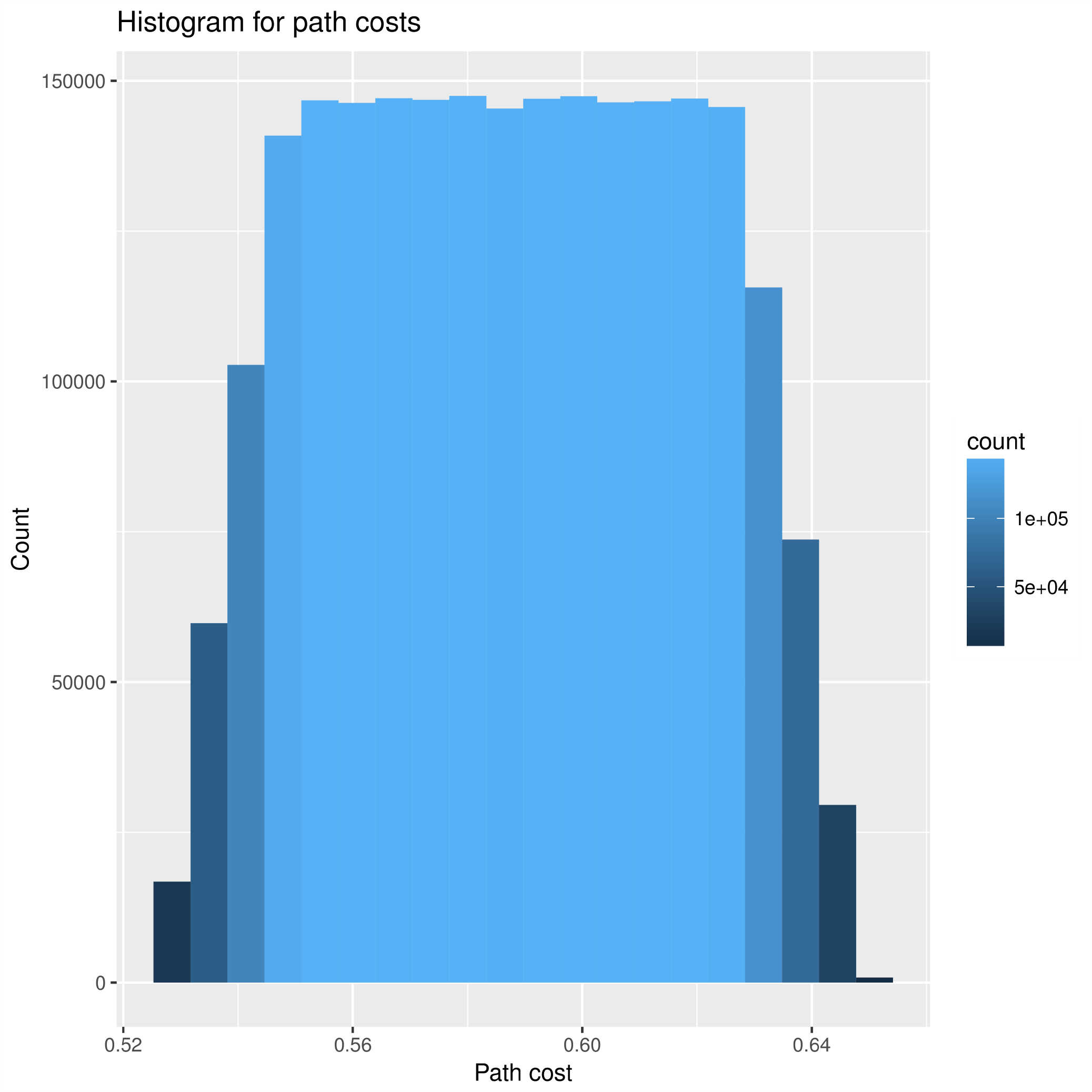
Path costs distribution for Infection Prevalence model. The shortest path is ‘anc-chimp ⇒ P. boisei ⇒ H. erectus’ with an average path cost 0.587.

#### HSV2-IT model

In this analysis, the input parameters of the gamma distribution: the proximity of species and the time-period overlap between species was varied by 10%. The distribution of individual path costs used to populate the edge costs of the DAG used in the sensitivity analysis are shown in fig. 5. Sensitivity analysis predicted a primary transmission route (60% of the cases) of ‘anc-chimps → H. habilis → P. boisei → H. erectus‘ and a secondary transmission route (remaining 40%) through ‘anc-chimps → P. boisei → H. erectus‘. The distribution of the shortest path costs for both the primary and secondary transmission routes are shown in fig. 6. The Sobol indices for the HSV2-IT model are presented in table 2.

**Figure 5.**
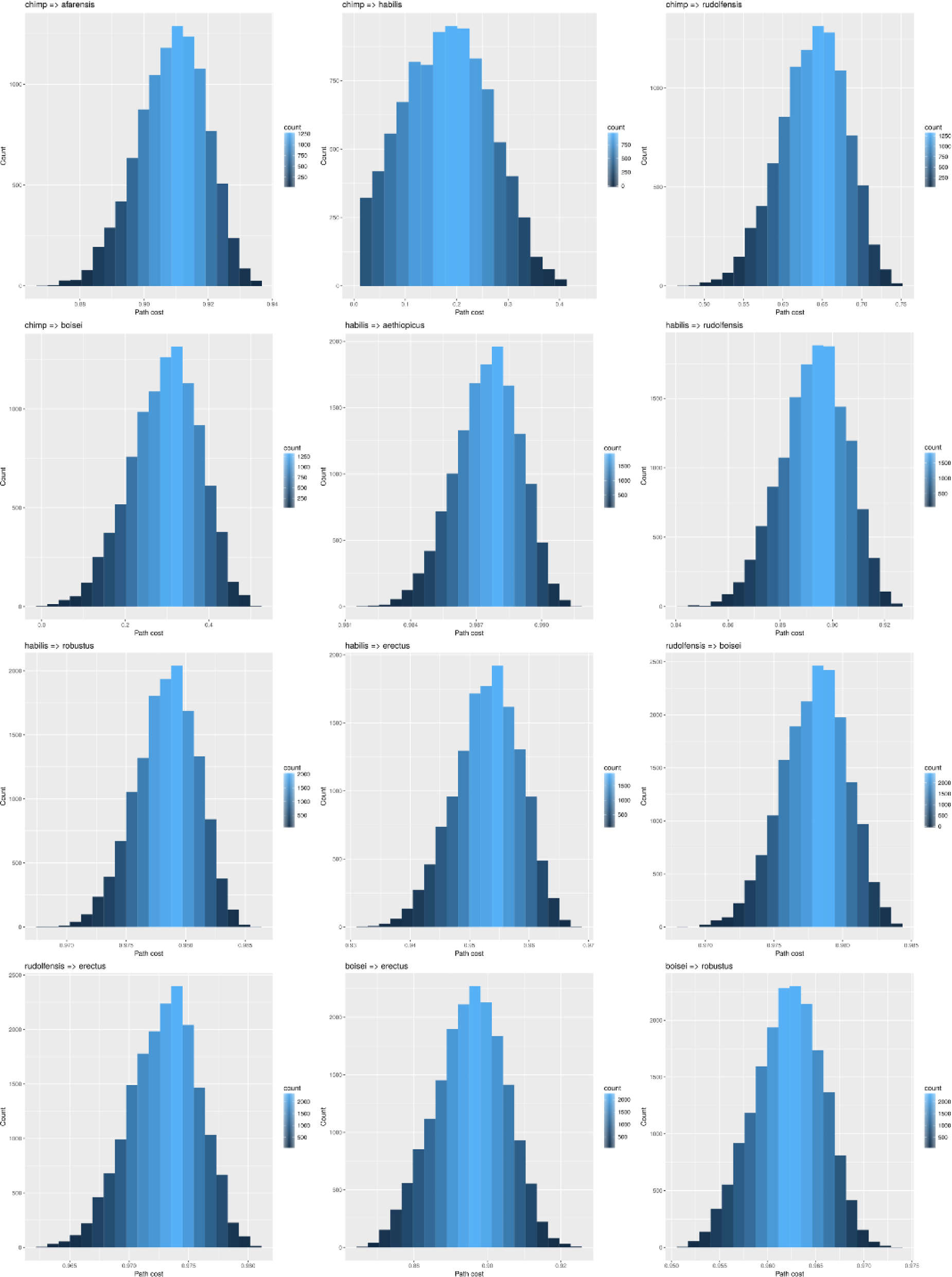
Distribution of selected edge costs for Infection Transmission model. Each histogram shows the distribution of edge costs for the IT model of HSV2 transmission. Colour contours indicate the number of occurrences of a path cost in the sensitivity analysis. The lowest edge costs are seen for ‘anc-chimp ⇒ H. habilis’, ‘anc-chimp ⇒ P. boisei’ and ‘P. boisei ⇒ H. erectus’.

**Figure 6.**
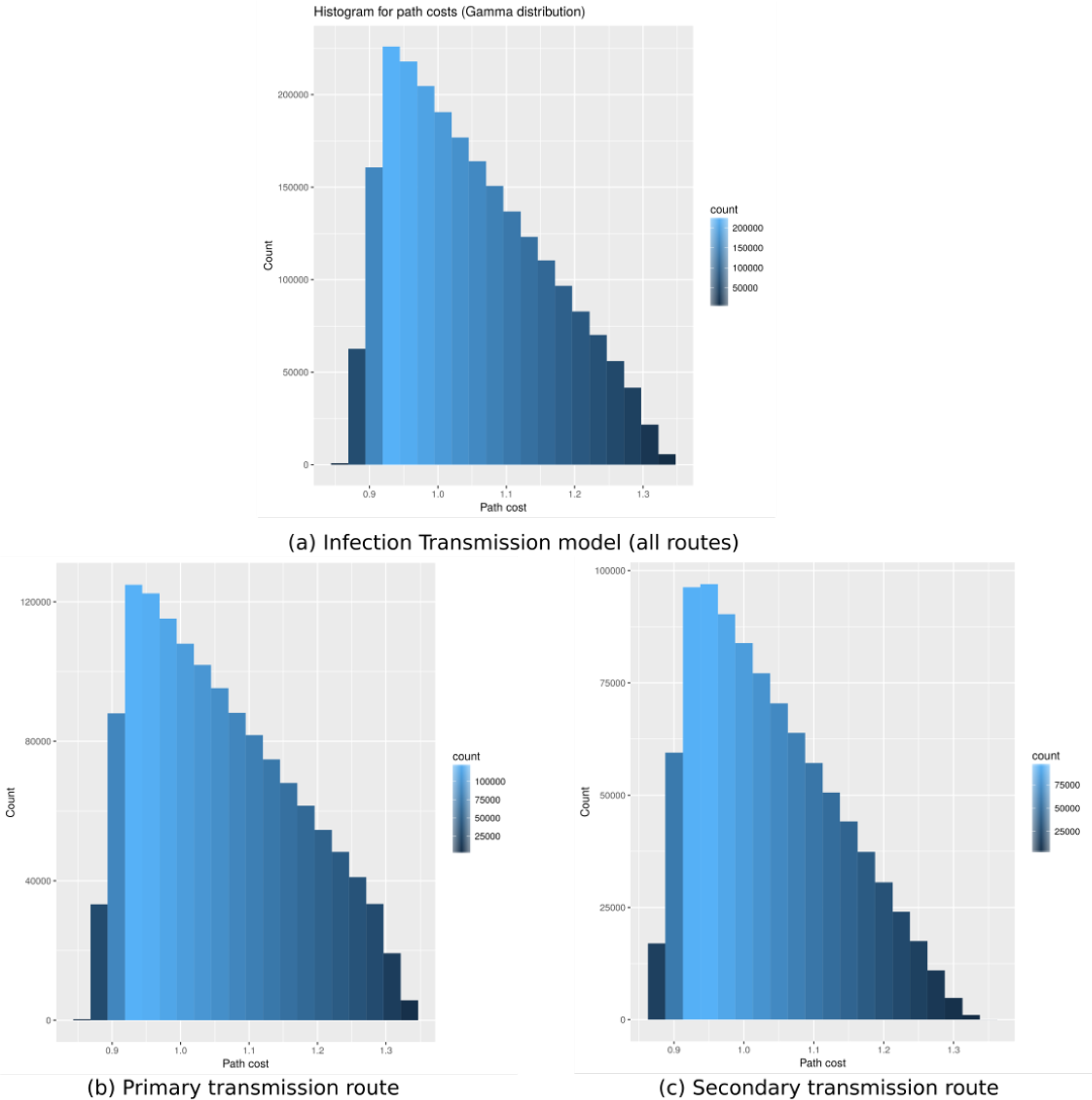
Path costs distribution for Infection Transmission model. Each histogram shows the distribution of path costs for the IT model of HSV2 transmission. Colour contours indicate the number of occurrences of a path cost in the sensitivity analysis. The optimal transmission route is ‘anc-chimp ⇒ *H. habilis ⇒ P boisei* ⇒ *H. erectus’.* (a) Total distribution of path costs for HSV2-IT model, (b) distribution path costs for the primary transmission route (‘anc-chimp ⇒ *H. habilis ⇒ P boisei* ⇒ *H. erectus’),* and (c) distribution path costs for the secondary transmission route (‘anc-chimp ⇒ *P boisei* ⇒ *H. erectus’).*

**Table 2.** Sobol indices of transmission paths for HSV2-IT model using gamma distribution.

| Disease Transmission Route | Sobol total-order indices | Sobol first-order indices |
| --- | --- | --- |
| anc-chimp ⇒ *P. boisei* | 0.640 | 0.454 |
| anc-chimp ⇒ *H. habilis* | 0.519 | 0.334 |
| *P. boisei* ⇒ *H. erectus* | 0.0262 | 0.0262 |

Fossils from four genera (Ardipithecus, Kenyanthropus, Orrorin and Sahelanthropus) were excluded from the analysis on the basis that there is no fossil evidence that they persisted after 3 MYA. Our range of candidate species can be restricted to those in supplementary Table A.1.

We then used further data on geo-temporal proximity of one species to another to develop mathematical models to quantify the probability of HSV2 infection transmission so as to assess the likelihood that each of these species further transmitted HSV2 to another hominin (see methodology section for further detail of the model). If HSV2 was transmitted to *Homo erectus,* parsimony militates against the need for a further cross-species transmission event to be invoked to explain the infection of modern humans by HSV2. Simple vertical mother-to-child or horizontal (sexual) transmission of the virus through the genus Homo from this point would be sufficient as the ancestor-descendent path from *Homo erectus* to *Homo sapiens* is relatively secure (Maslin, Shultz, and Trauth, 2015).

### Discussion

#### Transmission of HSV2

Transmission of HSV2 The combined inference analysis of a Bayesian Network, considering the presence of HSV2 in both anc-chimps and Homo erectus, revealed *A. afarensis* (90%), *H. habilis* (67.8%), *P boisei* (50%), and *H. rudolfensis* (55.8%) to be the likely intermediary hosts. Although, *A. afarensis* had a close proximity to the geographical range of anc-chimps, it could only transmit HSV2 to *A. africanus.* However, *A. africanus* has a mere 3% chance of being infected by HSV2 due its geographic location. This limits the possibility of HSV2 transmission to *H. habilis, P. boisei* and *H. rudolfensis,* either directly or as intermediary hosts.

The Infection Prevalence (HSV2-IP) is a local model that describes the probability of HSV2 infection based on the proximity to the rainforest habitat and the duration a species persisted in the fossil record using a beta distribution. In a beta distribution, the parameters alpha (time period) and beta (proximity to the rainforest) are weighted equally, i.e., an increase in the distance to the rainforest by 1 kilometer has the same effect on probability as a decrease of 1000 years in the time period. Thus the beta distribution assumes a constant rate of infection transmission. The HSV2-IP model considers each transmission as an independent event, i.e., the probability of a species transmitting HSV2 is independent of previous events leading to the HSV2 infection of that species.

The A* algorithm using the weighted DAG with the HSV2-IP model predicted the transmission of HSV2 from anc-chimps to *H. erectus* through *P boisei* with an average path cost of 0.587. Sensitivity analysis did not show any variation in the route (i.e., varying the location and the time-period of the probable species by 10% did not affect the optimal route). Non-zero values of sensitivity indices were observed only along the critical transmission route of ‘anc-chimp → P. boisei → *H. erectus’.* Although, the ‘P *boisei → H. erectus’* path accounted for 95.5% variation in the path costs, the path cost of ‘anc-chimp → *P. boisei*’ was the critical value that controlled the transmission route. Hence, the optimal route through *P. boisei* remained unaffected. Further analysis of the route revealed a significant influence (88 - 90% of the variation in the results) of the proximity and duration of *P boisei* in contrast to the probability of anc-chimps and *H. erectus* on the path costs. The infection prevalence model uses a beta distribution, which assumes equal weights for the shape parameters: *alpha* and *beta.* As expected, Sobol analysis showed equal influence for the distance and time-period parameters (a first-order Sobol indices of 0.493 and 0.492 respectively) on the path costs. Due to the nature of beta distribution, care should be taken in defining the rate of transmission (ratio of the shape parameters, alpha and beta), as the probability value estimated from the distribution is dependent on the ratio of the shape parameters. The intermediary host identified as P *boisei* is only 35 kilometers from the rainforest, which results in high probability of transmission, and hence results in the lowest cost of transmission through this species. HSV2-IP is a local model, which is independent of other events and does not consider the history of transmission and the associated probabilities.

The Infection Transmission (HSV2-IT) model assigns probability values (edge costs) to the DAG utilising the temporal overlap between hominin species and their geographic proximity to one another. The HSV2-IT model is history dependent, i.e., the probability of a child node transmitting a disease depends on the probability of the parent nodes infecting the child node in a DAG. The Infection Transmission model predicts an initial transmission of HSV2 from anc-chimps to *H. habilis.* Further, *H. habilis* transmitted the virus to *P boisei,* which then infected *H. erectus.* The average shortest path cost was estimated as 1.05. Sensitivity analysis of the HSV2-IT model revealed two transmission routes: (a) Primary transmission route (60% of the cases) of ‘anc-chimps → *H. habilis* → *P boisei* → *H. erectus’* and (b) a secondary transmission route (remaining 40% of the cases) through ‘anc-chimps → *P boisei* → *H. erectus*’. However, the distribution of path costs through both the routes were similar, showing that both transmission routes could have been possible. Sensitivity analysis shows that the ‘anc-chimp → *H. habilis’* and ‘anc-chimp → *P boisei’* have quite a significant effect on the transmission route (total path cost). About 70% the variation in the route of ‘anc-chimp → *H. habilis’* route is due to the direct effect of edge weight determined by the gamma distribution in comparison to 64% of variation for ‘anc-chimp → P. boisei’ route. Secondary interactions (interactions between different paths) accounts for 30 - 35% of the overall routing. The probability of a species transmitting, and another being infected had similar effects (similar Sobol indices), which explains that the model did not have a bias either toward the proximity or the temporal overlap between species. Because the HSV2-IT model considers the history of transmission, it is a better predictor than the HSV2-IP model.

Our analysis suggests that *Paranthropus boisei* was the most critical intermediate host for transmitting HSV2 between anc-chimps and the ancestors of *Homo sapiens.* The transmission of proto-HSV2 from anc-chimps to *P. boisei* could have happened directly or through *Homo habilis.* However, our analysis did not predict *Homo habilis* transmitting HSV2 directly to *Homo erectus.* Our results suggest the initial transmission to *P boisei* could have happened from *H. habilis* 60% of the time due (most likely through direct sexual contact rather than consumption of infected tissue) and the remaining 40% was directly transmitted from anc-chimps via hunting or scavenging.

*P. boisei* would have been well placed to act as an intermediate host for HSV2. It most likely contracted the infection through hunting or more likely scavenging infected ancestral-chimpanzee meat. Processing (with or without tools) and consumption of raw meat would act as a simple route for ChHV1 to have crossed into *P. boisei* via open cuts or sores. Tropical refugia during hot dry periods may have driven chimpanzees into higher concentrations in certain areas, driving them into contact and competition with P *boisei* and *H. habilis* as the margins of tropical forest blended into more open savannah like habitats (Julier et al., 2017). Violent confrontation or hunting/scavenging and butchery practices would have provided a viable route of transmission for HSV2. Homo habilis remains have been recovered from the same layers as stone tools and bones carrying evidence of butchery, supporting a possible transmission-through-hunting/scavenging hypothesis for the initial anc-chimp to *H. habilis* transmission (Clarke, 2012). *Paranthropus aethiopicus, P boisei,* and *P robustus* are associated with the Oldowan stone tool complex (De Heinzelin et al., 1999), and *P boisei* explicitly with butchery (Dominguez-Rodrigo et al., 2013) lending support to the hypothesis that bushmeat hunting/scavenging and butchery may have led to the initial transmission of HSV2 to the hominins.

Both *Homo erectus* and *Paranthropus boisei* are known from sites around Lake Turkana in Kenya that are contiguous in age (Anton et al., 2016; Wood and Constantino, 2007) and it is likely that close contact between the species would have been relatively common especially around water sources. The appearance of *Homo erectus* 2.0 MYA is accompanied by evidence of active hunting and butchery, and from 1.76 MYA increasingly sophisticated stone tools (Cachel and Harris, 1998). This behavioural shift towards active hunting displayed by *Homo erectus,* combined with direct archaeological evidence of hunting, provides a credible route for direct transmission of HSV2 from *P boisei* to *H. erectus* through contact and/or consumption of infected material processed from *P boisei* carcasses.

In this study, *Homo erectus* is chosen as the target node for the transmission of HSV2 for a number of reasons. Morphological adaptation for bipedal locomotion is used as the primary trait for assigning fossils to the hominin sub-family, especially between 7.0 - 4.5 MYA, but is not an effective tool for determining patterns of ancestor-descendant relationships in the fossil record. The hominin sub-family currently contains seven genera but the exact taxonomic relationship between each genus is not clearly definable because of the fragmentary nature of the fossil record. Similarly, patterns of intra- and inter-species variation are difficult to define (Robert A Foley, 2016). The adaptive radiation of the genus *Australopithecus* between 4.5 - 2.0 MYA represents the first morphologically coherent group of fossil hominin species but its relationship to the genus Homo is not yet clear. The appearance of *Homo erectus* circa 2.0 MYA in East Africa marks the first appearance of recognisably ’human’ morphology, life history and brain development and represents a secure most recent common ancestor (MRCA) for all subsequent Homo species (with the possible exception of *Homo floresiensis* (Argue et al., 2017; Anton et al., 2016). Therefore, the ancestor-descendant relationship between *Homo erectus* at c. 2.0 MYA and *Homo sapiens* c. 200 KYA is an evolutionary secure route of transmission for HSV2 to leave Africa as a modern human-borne virus.

Although ChHV1 causes outbreaks of oral and pharyngeal lesions in chimpanzees in a manner similar to HSV1, in hominins contracting proto-HSV2, the oral niche was already occupied by HSV1. This may have protected the hominin first infected with HSV2: pre-existing infection with HSV1 reduces the likelihood that subsequent infection with HSV2 will be symptomatic (Langenberg et al., 1999), and also reduces the risk of HSV2 meningitis (Aurelius et al., 2012). HSV2 may have been forced to adapt to a different mucosal niche in order to reduce competition from the co-evolved, native HSV1. However, both simplexviruses remain capable of infecting both the oral and genital niche in modern humans (Kim et al., 2006; Whitley, Kimberlin, and Prober, 2007), and of causing co-infection in both niches (eg 10% to 15% of herpes labialis (oral lesions) is caused by HSV-2 (Glick and Siegel, 1999)).

We suggest that the mode of transmission of HSV2 to hominins was most likely through hunting injuries (eg chimpanzee bites or cuts sustained during meat processing), although onwards transmission into the ancestors of *Homo sapiens* could have been sexual (horizontal) or a result of hunting injuries (vertical). There are many reports of transmission of B virus *(Cercopithecine herpesvirus* 1), the cercopith homolog of HSV1 and ChHV1, to humans, where disease ranges from mild to fatal. Transmission has occurred from bites and scratches, needle sticks and even scratches from cage bars that are contaminated with B virus-positive bodily fluids (Huff and Barry, 2003). Onwards transmission between humans has been reported to occur (Disease Control, Prevention, et al., 1987). HSV1 can infect other primates, from gorillas (Gilardi et al., 2014) to owl monkeys (Melendez et al., 1969), typically causing fatal disease in species more distantly related to *Homo sapiens,* while causing oral lesions and milder disease in great apes such as *Gorilla beringei graueri* (Gilardi et al., 2014). We therefore infer that herpes simplex-like viruses spread relatively easily between individuals even across species barriers, increasing the chances of transmission between hominins and other primates from close contact such as hunting, butchery, inter-personal violence or sexual contact. Evidence for close hominin-hominid contact is also found in other *‘heirloom’* human pathogens (Houldcroft et al., 2017).

The high prevalence of HSV2 in central and eastern Africa (Looker et al., 2015) (Supplementary Fig.1 and interactive maps at https://wadhamite.github.io/hsv2-map) is consistent with the limited genetic data available from African HSV2 isolates. A study from Burrel and colleagues (Burrel, Desire, et al., 2015) showed that HSV2 can be divided into African and worldwide lineages on the basis of diversity in gene UL30. Furthermore, two groups found evidence of gene flow from HSV1 into HSV2, and speculated that the flow of HSV1 loci into the worldwide HSV2 lineage may have helped this lineage of HSV2 to further adapt to human hosts, and so spread more successfully around the world from around 41 KYA (Burrel, Desire, et al., 2015; Koelle et al., 2017). Recent studies have significantly increased the number of whole HSV2 genomes available for analysis, contributing to our knowledge of HSV2 diversity (Kolb et al., 2015; Szpara et al., 2014); but there is conflicting evidence on whether the most basal HSV-2 genotypes are from west and central (Burrel, Boutolleau, et al., 2017) or east (Koelle et al., 2017) Africa. Our analyses predict that individuals from east Africa are likely to carry the most ancient HSV2 lineages.

The time-depth of ancient DNA analysis is dually limited by technology and preservation of DNA. Similarly, the archaeological and fossil records suffer from differential rates of preservation (Allentoft et al., 2012) and gaps that can never be filled because the material has simply not survived. Our analysis has allowed the reconstruction of hominin/human-disease interaction well beyond the horizon of ancient DNA and at a level that is invisible to the fossil and archaeological records. There are other ancient human pathogens that have switched between different primate and hominin hosts over the last 6 million years and these transmission routes could be further explored with this methodology. For example, human pubic lice *(Pthirus pubis)* were introduced by an unknown hominin through contact with the ancestor of gorillas around 3.3 MYA (D. L. Reed et al., 2007). There is also evidence of Neanderthal to human transmission of human papillomavirus genotypes (Pimenoff, Oliveira, and Bravo, 2017), and of hominin to human transmission of body louse genotypes (D. D. L. Reed et al., 2004). The studies and the findings presented here demonstrate the potential for using modern disease genetics to understand better the evolutionary interaction between humans and disease in deep time.

## Data presentation

The distribution maps were created with custom JavaScript codes using Leaflet.js library [http://leafletjs.com] and interactive maps can be accessed at https://wadhamite.github.io/hsv2-map

## Author contributions

SJU and CH conceived the study and contributed data. SJU, KK and CH performed the analyses. SJU, KK and CH wrote the paper. All authors approved the publication of the manuscript.

## Acknowledgements

The authors would like to thank C Ruis (University College London), and JB Ramond and RF Rifkin (University of Pretoria) for helpful discussion. SJU was funded by Oxford Brookes University. KK and CH were funded by the University of Cambridge. KK is a college research associate at King’s College, Cambridge. CH is a post-doctoral affiliate at Christ’s College, Cambridge.

## Supplementary material

**Figure A.1.**
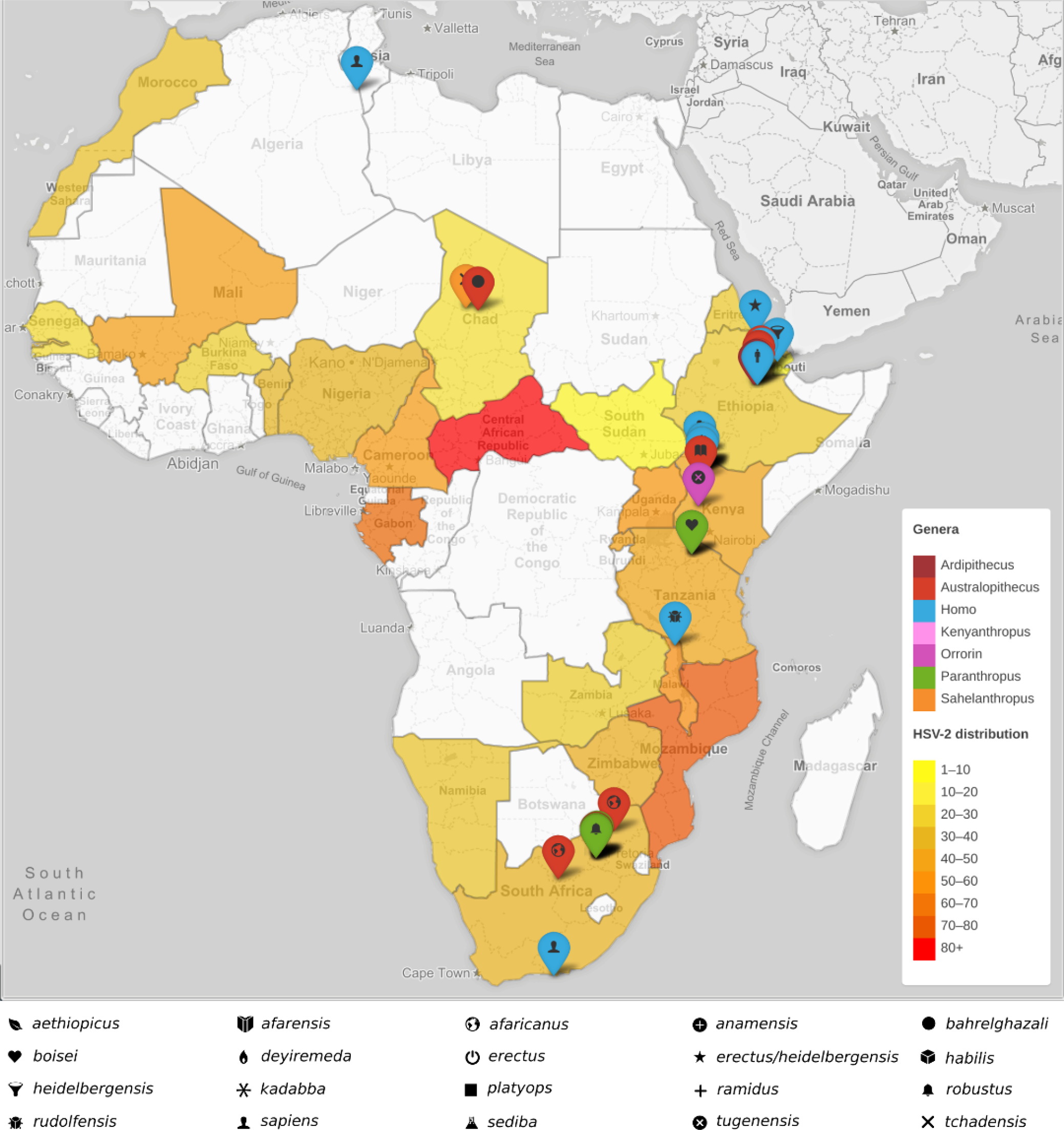
Map showing the prevalence of HSV2 in Africa, using data from Looker (2015). The locations of hominin fossils [supplementary table 1] are shown with markers. The colour of the marker indicates the hominin genus; the symbol represents the species. This figure is available interactively: https://wadhamite.github.io/hsv2-map

**Table A.1.**
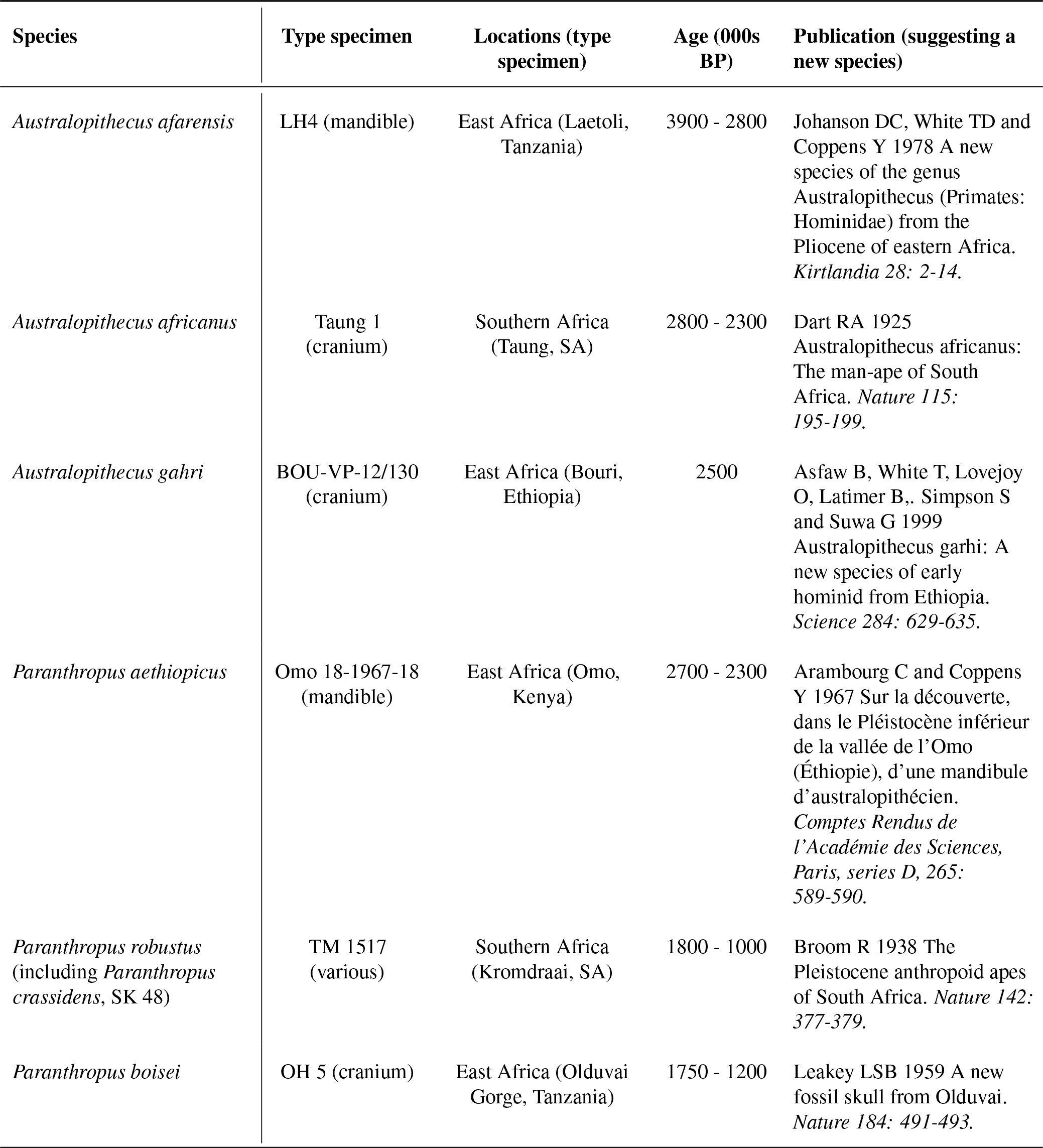
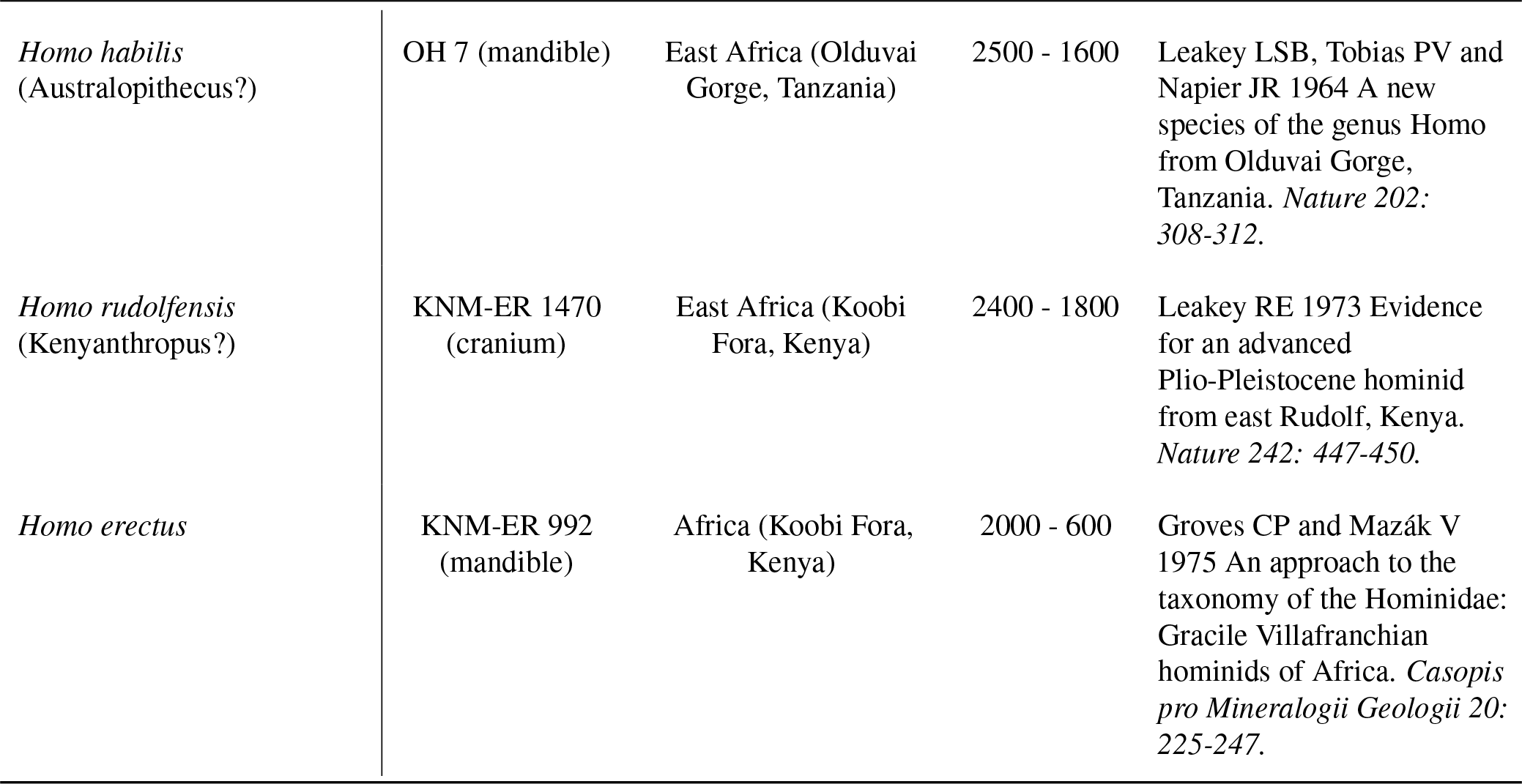
Fossil hominins with the potential to acquire - HSV2 from ancestral chimpanzee

| Species | Type specimen | Locations (type specimen) | Age (000s BP) | Publication (suggesting a new species) |
| --- | --- | --- | --- | --- |
| *Australopithecus afarensis* | LH4 (mandible) | East Africa (Laetoli, Tanzania) | 3900 - 2800 | Johanson DC, White TD and Coppens Y 1978 A new species of the genus *Australopithecus* (Primates: Hominidae) from the Pliocene of eastern Africa. *Kirtlandia* 28: 2-14. |
| *Australopithecus africanus* | Taung 1 (cranium) | Southern Africa (Taung, SA) | 2800 - 2300 | Dart RA 1925 *Australopithecus africanus*: The man-ape of South Africa. *Nature* 115: 195-199. |
| *Australopithecus gahri* | BOU-VP-12/130 (cranium) | East Africa (Bouri, Ethiopia) | 2500 | Asfaw B, White T, Lovejoy O, Latimer B., Simpson S and Suwa G 1999 *Australopithecus garhi*: A new species of early hominid from Ethiopia. *Science* 284: 629-635. |
| *Paranthropus aethiopicus* | Omo 18-1967-18 (mandible) | East Africa (Omo, Kenya) | 2700 - 2300 | Arambourg C and Coppens Y 1967 Sur la découverte, dans le Pléistocène inférieur de la vallée de l'Omo (Éthiopie), d'une mandibule d'australopithécien. *Comptes Rendus de l'Académie des Sciences, Paris, series D*, 265: 589-590. |
| *Paranthropus robustus* (including *Paranthropus crassidens*, SK 48) | TM 1517 (various) | Southern Africa (Kromdraai, SA) | 1800 - 1000 | Broom R 1938 The Pleistocene anthropoid apes of South Africa. *Nature* 142: 377-379. |
| *Paranthropus boisei* | OH 5 (cranium) | East Africa (Olduvai Gorge, Tanzania) | 1750 - 1200 | Leakey LSB 1959 A new fossil skull from Olduvai. *Nature* 184: 491-493. |
| *Homo habilis*<br>(Australopithecus?) | OH 7 (mandible) | East Africa (Olduvai Gorge, Tanzania) | 2500 - 1600 | Leakey LSB, Tobias PV and Napier JR 1964 A new species of the genus *Homo* from Olduvai Gorge, Tanzania. *Nature* 202: 308-312. |
| *Homo rudolfensis*<br>(Kenyanthropus?) | KNM-ER 1470 (cranium) | East Africa (Koobi Fora, Kenya) | 2400 - 1800 | Leakey RE 1973 Evidence for an advanced Plio-Pleistocene hominid from east Rudolf, Kenya. *Nature* 242: 447-450. |
| *Homo erectus* | KNM-ER 992 (mandible) | Africa (Koobi Fora, Kenya) | 2000 - 600 | Groves CP and Mazák V 1975 An approach to the taxonomy of the Hominidae: Gracile Villafranchian hominids of Africa. *Casopis pro Mineralogii Geologii* 20: 225-247. |

